# Skilful decadal-scale prediction of fish habitat and distribution shifts

**DOI:** 10.1101/2021.07.07.451446

**Authors:** Mark R. Payne, Anna K. Miesner, Noel Keenlyside, Shuting Yang, Stephen G. Yeager, Gokhan Danabasoglu, Daniela Matei

## Abstract

Many fish and marine organisms are responding to our planet’s changing climate by shifting their distribution (i.e. where they are found). Such shifts can drive conflicts at the international scale and are highly problematic for the communities and businesses that depend on these living marine resources for income and nutrition. Advances in climate prediction mean that in some regions the state of the ocean, and thereby the drivers of these shifts, can be skilfully forecast up to a decade ahead. However, the potential for these forecasts to benefit ocean-dependent communities has yet to be realised. Here we show for the first time that marine climate predictions can be used to generate decadal-scale forecasts of shifts in the habitat and distribution of marine fish species, as exemplified by Atlantic mackerel, bluefin tuna and blue whiting. We show statistically significant forecast skill of individual years that outperform both persistence and climatological baseline forecasts for lead times of 3-10 years: multi-year averages perform even better, yielding correlation coefficients in excess of 0.90 in some cases. We also show that the habitat shifts underling recent conflicts over Atlantic mackerel fishing rights could have been foreseen on similar timescales. Our results show that climate predictions can be translated into information directly relevant to stakeholders and we anticipate that this tool will be critical in foreseeing, adapting to and coping with the challenges of a changing and variable future climate, particularly in the most ocean-dependent nations and communities.

---

Our current understanding of the impacts of climate change typically focuses on the “climatic” time scale, i.e., 50 or 100 years into the future. While these timescales are of strategic value to governments and large international organisations, they are far from the seasonal, annual and decadal timescales on which regional bodies, local-governments, businesses and individuals make most of their decisions (Bruno Soares *et al.*, 2018). The recent development of near-term climate predictions (Kirtman *et al.*, 2013; Meehl *et al.*, 2014; Merryfield *et al.*, 2020) can potentially fill this gap and examples of such “climate services” can already be found on the sub-seasonal and seasonal timescales, primarily in terrestrial settings (Hewitt *et al.*, 2012; Street, 2016; Buontempo and Hewitt, 2018). Progress on the key annual-to-decadal timescales, where many strategic decisions are made, has been limited however and only the ocean currently has sufficient predictability (Yeager and Robson, 2017; Smith *et al.*, 2020) to support such forecasts. Nevertheless, the high ocean-dependency, vulnerability and climate risk of many coastal communities and countries, particularly in the Global South (Barange *et al.*, 2014; Golden *et al.*, 2016; Blasiak *et al.*, 2017), creates a pressing need for decadal-scale forecasts to support climate adaptation and sustainable development (IPCC, 2019).

Realising the potential of oceanic decadal-predictability, however, requires converting climate prediction data into information that addresses the challenges directly faced by stakeholders. One such challenge, and one of the most commonly reported impacts of climate change in the ocean, is shifts in where species are found (i.e., their “distribution”). Distributional shifts have been reported from the lowest trophic levels to fish and top-predators (Poloczanska *et al.*, 2016) and are occurring faster than on land due to the higher vulnerability of marine species to warming (Pinsky *et al.*, 2019). Projections suggest that this trend will continue with impacts being felt globally (Barange *et al.*, 2014; Blasiak *et al.*, 2017; Pinsky *et al.*, 2018; IPCC, 2019). As traditionally fished species disappear and new species arrive, local communities and fishers are required to adapt their fishing techniques, infrastructure, markets and even culinary preferences to the changed fishing opportunities. International conflicts over fishing rights can also arise as shifting fish stocks start to straddle international jurisdictions: transboundary stocks may impact as many as 40% of exclusive economic zones in the future (Pinsky *et al.*, 2018). Examples of such conflicts are already being seen (Spijkers *et al.*, 2019) (e.g., the so-called North Atlantic “mackerel war” (Spijkers and Boonstra, 2017) between the European Union, Norway, Iceland and the Faroe Islands over access to Atlantic mackerel, *Scomber scombrus*) and are a leading cause of international disputes between democracies (Mitchell and Prins, 1999; Spijkers *et al.*, 2019). The ability to foresee such shifts can therefore potentially hold the key to both avoiding conflict and adapting marine fisheries to a changing and variable climate.

Here we demonstrate the ability to forecast shifts in the habitat and distributions of marine species on the decadal scale for the first time. We focus on three exemplar fish species in the North Atlantic that have shown well-documented distribution shifts in recent years. The Northeast Atlantic stock of mackerel supports one of the most valuable fisheries in Europe and recent distribution shifts into Icelandic and Greenlandic waters (Jansen *et al.*, 2016) have driven the aforementioned conflict over fishing rights. Atlantic bluefin tuna (*Thunnus thynnus*) is a large commercially valuable and endangered top-predator: in recent years the species has shifted into the Irminger Sea and Denmark Strait (MacKenzie *et al.*, 2014), opening up new fishing opportunities for Iceland and Greenland (Jansen *et al.*, 2020). Blue whiting (*Micromesistius poutassou*) has at times been one of the world’s largest fisheries and its spawning distribution shifts regularly between the waters of the UK, Ireland, Faroe Islands and areas beyond national jurisdiction (Miesner and Payne, 2018), a potential problem in light of the UK’s departure from the EU. For each of these species we combined habitat models (MacKenzie *et al.*, 2014; Jansen *et al.*, 2016; Miesner and Payne, 2018) (Extended Data Table 1) with existing climate prediction systems (Extended Data Table 2) to produce decadal-scale habitat predictions and verified these predictions against habitat estimated from ocean observations.

We first show the ability to skilfully forecast the physical drivers that serve as inputs to the habitat models (see Methods). Sea surface temperature (SST) in the warmest month (August) is used in the mackerel and bluefin tuna predictions while sub-surface salinity (250-600 m) during the peak spawning month (March) is the primary environmental driver shaping blue whiting habitat. Predictive skill of these variables is generally high and statistically greater than zero in most parts of the domain (Fig.1), in line with more general results (e.g., annual averages) reported elsewhere (Yeager and Robson, 2017; Kushnir *et al.*, 2019). The skill also matches well with the regions relevant to each of the marine species that we consider, providing a solid base from which to develop ecological forecasts. Similar results are seen when considering the absolute error in the forecasts (Ext. Data Fig.1) rather than the correlation (Fig.1).

**Fig.1.**
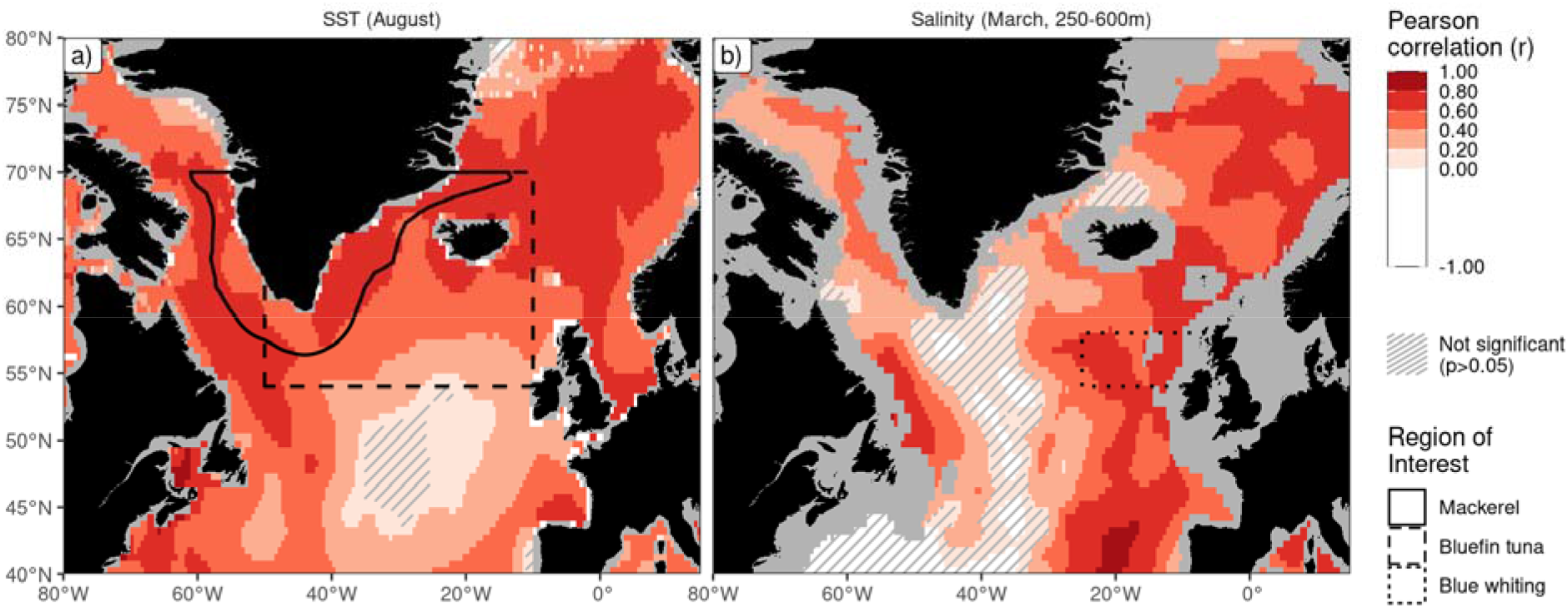
Physical forecast skill. Predictive skill of physical variables underlying ecological forecasts with a lead time of five years for a) mean August sea surface temperature (SST) and b) mean March sub-surface (250-600 m) salinity. Predictive skill is expressed as the Pearson correlation coefficient (*r*) between the forecast and observed values of each variable, with each grid point coloured according to the local value. Forecast skill is for the grand ensemble mean forecast, i.e., a forecast averaged across the individual realisations from all model systems. Regions where the correlation coefficient is not greater than 0 (at the 95% confidence level) are cross-hatched. Lines mark the polygons over which ecological forecasts are integrated in subsequent analyses. Ocean regions not represented by the models are shown in grey.

When outputs from the climate prediction systems are used in the habitat models, we see retrospective-forecast skill on both multi-annual and decadal time scales. We first integrate forecast maps of habitat over the relevant regions of interest to produce metrics of habitat area. Pearson correlation coefficients between the forecast habitat indicators and those derived from observations are generally high, up to 0.75 for the forecast including all ensemble members (“Grand ensemble”, Fig. 2). This skill remains high even at decadal lead times, and is significantly greater than zero (p < 0.05) for all leads and species. Individual modelling systems can have lower performance, but the combination of models into a grand ensemble generally gives the best performance.

**Fig. 2.**
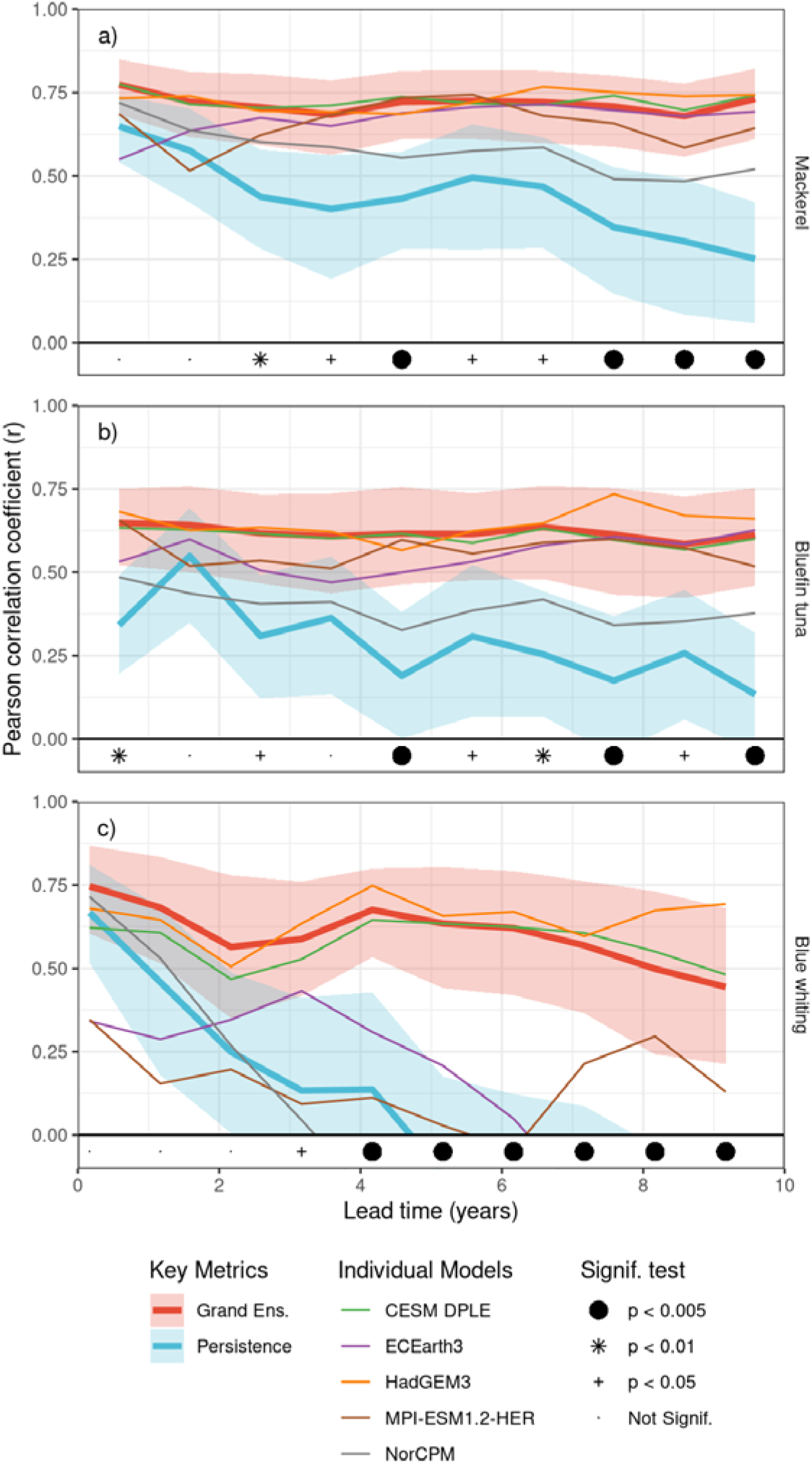
Habitat forecast skill. Skill for indicator metrics of a) mackerel, b) bluefin tuna, and c) blue whiting habitat. Forecast skill is given as the Pearson correlation coefficient (r) between the forecast habitat area and that derived from observational data, and is plotted as a function of forecast lead time into the future. Forecast skill is shown for the mean forecasts of the individual models (light weighted lines) and for the grand-ensemble forecast across all ensemble members (heavy red line). The skill of persistence forecasts (heavy blue lines) are also shown for reference. Shaded areas for both these key metrics denote the 90% confidence interval: 5% of the distribution is therefore above and 5% below the shaded areas. The hypothesis that the grand-ensemble forecast outperforms persistence (i.e. a one-tailed test) is tested for each lead time, and denoted with symbols at the bottom of each panel.

Importantly, our ecological forecast systems also outperform alternative simpler approaches. We consider a persistence forecast, where “tomorrow is the same as today”, as a much simpler and commonly used baseline system: a valuable forecast system should have skill over and above this reference forecast (Joliffe and Stephenson, 2012). For short lead times (e.g., one-two years), persistence forecasts are generally comparable to climate predictions (Fig. 2), reflecting the high inertia of the ocean. In these cases, the improvement in skill of ecological forecasts over persistence forecasts is generally not significant or at best weak. On the multi-annual scale, however, persistence skill starts to fade while the decadal prediction systems maintain their forecast skill. For all three fish stocks considered, forecast performance for leads of three or more years is significantly greater than persistence (p < 0.05 or better), and can therefore be considered skilful. Alternative skill metrics considering the absolute errors in the forecasts (via the mean-squared error skill score) and reliability of the predictive distributions (continuous ranked probability skill score) also show significant skill across all fish stocks and for multi-annual to decadal lead times (Extended Data Fig. 2). As is common in multi-model ensemble systems (Palmer *et al.*, 2005), the grand-ensemble habitat forecast based on all 85 members weighted equally is also consistently the amongst the best performing forecasts (Fig. 2) reiterating the value of large ensembles in climate prediction (Smith *et al.*, 2020).

While we capture the majority of the variability when forecasting individual years, the forecasts perform even better when considering multiple years. We calculated multi-year means of habitat estimates derived from both the climate prediction systems and observational data, and then re-evaluated the forecast skill (Fig. 3a–c). Improvements in skill are seen for all fish stocks, reaching correlation coefficients of 0.95, and 0.74 for predictions of the decadal average (9 year window) for mackerel, bluefin tuna and blue whiting respectively. Averaging also improves some persistence forecasts but the ecological forecast system continues to be significantly better (p < 0.01 for 9 year averages across all stocks). The ultimate choice of averaging window clearly depends on the needs of the decision maker using the forecast: short-term tactical planning may need the individual years, while longer-term strategic planning may require the decadal averages or the statistical distribution. Importantly and reassuringly, we show significant decadal forecast skill of both the mean (pearson correlation, MSESS metrics) and the distribution (CRPSS metric), with and without averaging.

**Fig. 3.**
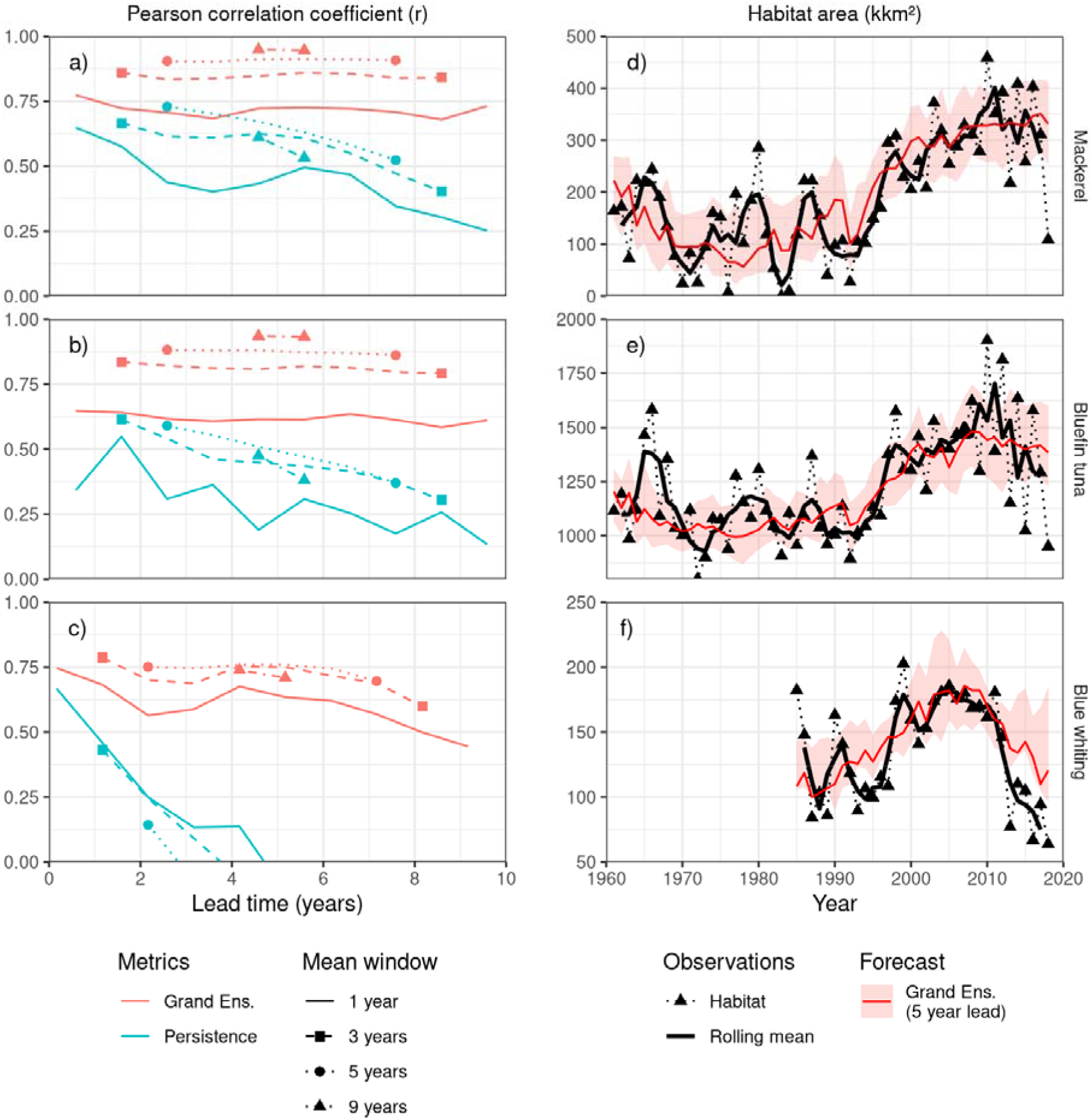
Sources of forecast skill. The skill of multi-annual forecasts (panels a-c), as characterised by the Pearson correlation coefficient (*r*), is shown for the grand-ensemble and persistence forecasts. In addition to the single-year values also plotted in Fig. 2 (solid lines), the skill of multi-year averages (3, 5, and 9 year centred means) are also shown (broken lines with symbols). Lead-time is defined as the length of time from the issuing of the forecast (1 January) to the middle of the running mean window. Multi-year forecasts are significantly better than multi-year persistence for all lead times (p < 0.01). Time-series of habitat indicators (panels d-f) show observations (triangles connected by dotted line) with their three-year running means mean (solid black lines). Habitat metrics forecast by the grand-ensemble (solid red line) with a 5-year lead time are shown with the corresponding 90% range of realizations (shaded area). Time series are shown for the full range of years used to estimate the forecast performance (i.e., 1961-2018 for mackerel and bluefin tuna, 1985-2018 for blue whiting). Panels a) and d) show results for the area of mackerel habitat around south Greenland, panels b) and e) bluefin tuna habitat south of Iceland, and c) and f) blue whiting spawning habitat west of Great Britain and Ireland.

The effect of multi-year averaging on our predictions is closely linked to the source of their skill. On short timescales, process originating from atmospheric dynamics (e.g. blocking highs) strongly influence oceanic variability, especially for SST: while these processes are present in climate models, they are not predictable beyond “weather” timescales due to their chaotic nature. On longer timescales, the North Atlantic sub-polar gyre, Atlantic Multidecadal Variability, and anthropogenic warming set the background oceanographic conditions (on top of which high-frequency variability is imposed): the representation of all of these aspects of Atlantic thermohaline circulation benefit from the initialisation process in climate prediction models and are well predicted (Smith *et al.*, 2019). The ability to capture the lower-frequency variability of the physical system also propagates into our ecological forecasts, which are clearly better at capturing multi-annual variability than interannual (e.g. Fig. 3d–f). Multi-annual averaging improves these forecasts further by filtering out the high-frequency interannual-noise, thereby increasing the relative contribution of the predictable low-frequency components. The skill of our ecological forecasts is therefore primarily due to the strong low-frequency (decadal) variability in the system, together with the ability of initialised climate prediction models to capture these slower processes.

The potential value of these forecasts to users can be illustrated by considering individual events. For example, the seas around Greenland supported very little (and at times no) water that was warm enough to act as mackerel habitat at the start of the 1990s (Fig. 4a). However, from 1990 to 2010, habitat availability of Greenlandic waters increased four-fold, ultimately facilitating the expansion of mackerel into this region (Jansen *et al.*, 2016). Decadal forecasts issued from 1990 onwards could have foreseen first the rapid expansion, and then its subsequent slowing after 2000. Decision makers deciding whether to pursue a commercial fishery after the first catches of Mackerel in Greenlandic waters in 2011 (Jansen *et al.*, 2016) could have been encouraged by decadal forecasts that (correctly) predicted that the expanded habitat would persist for the next decade. Decadal-scale ecological forecasts of habitat changes therefore clearly have a role to play in supporting strategic decision-making in the marine sector.

**Fig. 4.**
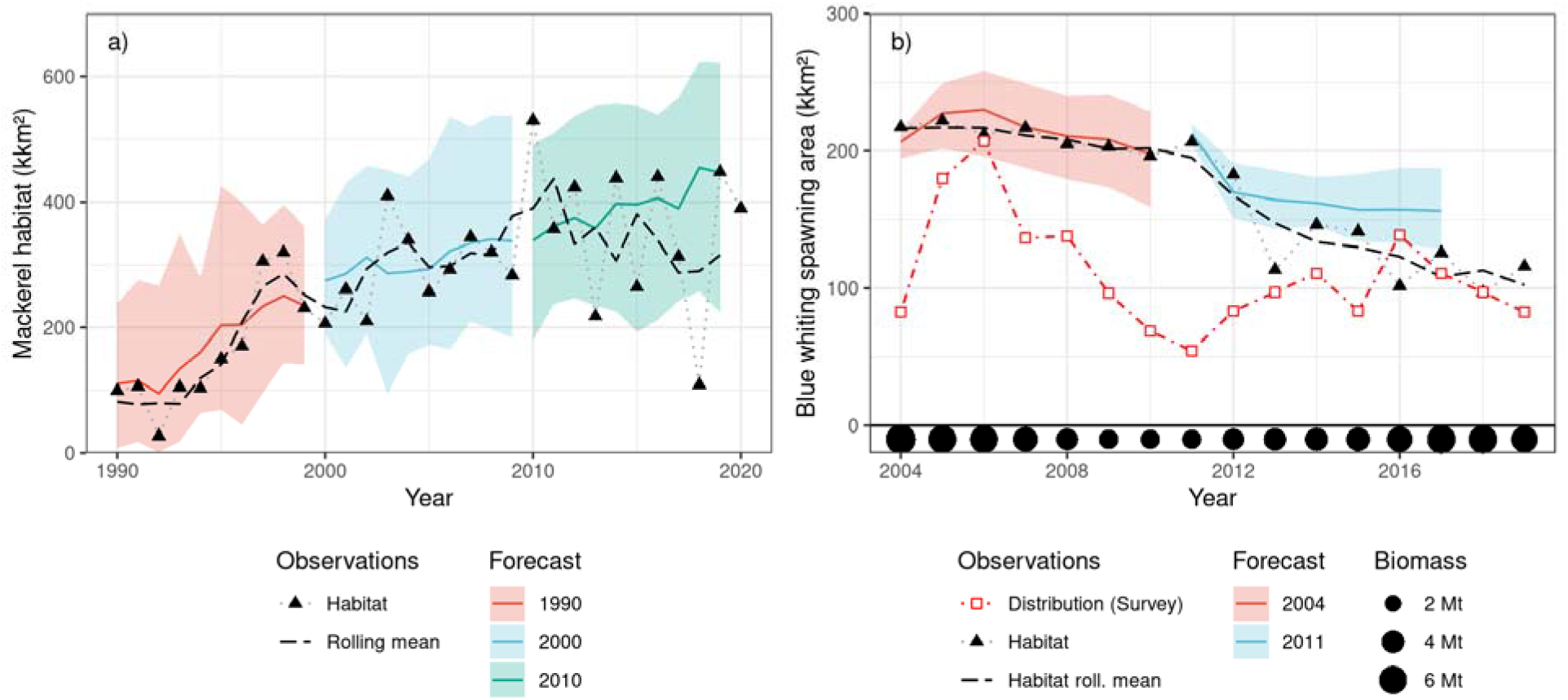
Forecasts of habitat and distribution changes. Habitat forecasts from the best performing CESM-DPLE model system started at select times (solid coloured lines) are shown with the corresponding 90% range of realizations in this model (shaded colours area) for the area of a) mackerel habitat around south Greenland and b) blue whiting spawning habitat. Habitat metrics based on observations (see Method) in a given year are shown (triangles connected by dotted grey line) together with a three-year centred running mean (dashed black line) of these values. For b), blue whiting, the distributional area estimated from scientific monitoring surveys (red dashed line) is shown together with the spawning stock biomass (bullets at bottom of figure) estimated by the stock-assessment. An illustrative subset of predictions is shown on both panels: the full set can be seen in Fig. 3d–f.

It is important to note that the ecological niche models used here represent the habitat of the individual species, and not their distribution. Habitat, in this context, corresponds to the range of potential spatial locations where the species could potentially be found, whereas distribution refers to where the species actually was (or will be) found. Many processes influence the way in which species do, or do not, use their available habitat, including competition, presence (or absence) of predators, schooling and migration, behavioural dynamics, and the need to close the life cycle (i.e. reproduce) (Guisan and Zimmermann, 2000). Furthermore, habitat is further constrained by environmental factors that are not included in these models (e.g. food quantity and quality). Forecasts of the presence of habitat should therefore be viewed as a necessary, but not sufficient, condition to observe a species at a given location: the presence of habitat does not guarantee the presence of fish. On the other hand, the absence of habitat does indeed guarantee the absence of fish. The real-world skill of predicting distribution shifts is therefore asymmetrical because these habitat models are much better at predicting absence than presence.

The recent decline in the spawning habitat of blue whiting illustrates this interplay clearly (Fig. 4b). In the mid-2000s, when the blue whiting stock was at its highest recorded level, the observed area of the distribution closely corresponded to habitat estimated from oceanographic observations. While the amount of this habitat slowly declined (a feature predicted by decadal forecast systems), the area of the distribution collapsed much more rapidly as the stock shrank due to a high fishing pressure. Recovery of the stock was accompanied by expansion of the distribution, but only back to 2/3 of the area seen earlier (as predicted by the forecast systems). Habitat forecasts can therefore be used to infer distributional changes in some instances.

The interpretation of these results is influenced by the baseline forecast used. Here we used “persistence” as our baseline, the simplest and most common “model” for many decision makers. An alternative approach common in the scientific community is to use “uninitialized” climate *projections* (not predictions) as the baseline (e.g. Matei *et al.*, 2012b). Making this comparison showed initialised forecasts to be consistently better than uninitialized forecasts for skill metrics that focus only on the mean value (i.e. Pearson correlation, MSESS), although the difference was only significant for blue whiting (p < 0.05 for lead times up to 8 years, Extended Data Fig. 3a, b). However, the ability of the initialised models to capture the probability distribution of habitat (as indicated by the CRPSS metric) was significantly better than the uninitialized forecasts (Extended Data Fig. 3c). This result is consistent with expectations (Kirtman *et al.*, 2013): initialisation pushes the predictions towards observations and narrows their distribution compared to uninitialized models, yielding forecasts that are both more accurate and more precise. Given the well-established need to communicate both the most likely value and the potential range of values (i.e. uncertainty) together in a forecast (Bruno Soares and Dessai, 2016), the better probabilistic performance of initialised systems makes them clearly preferable to uninitialized models in these cases.

While the ability to forecast habitat and distribution is potentially valuable to users, avoiding conflicts due to shifting distributions requires more than just reliable predictions. For example, stakeholders also need to have the ability to act on this information (Pfaff *et al.*, 1999; Bruno Soares *et al.*, 2018). Distribution shifts will often result in both “winners” and “losers” and there is therefore a natural tendency on the part of the negatively impacted party to resist change. International agreements for managing such transboundary stocks need to have sufficient flexibility to cope with distributional shifts, while at the same time ensuring the sustainability of both the agreement and the fish stock itself (Pinsky *et al.*, 2018). Decadal forecasts of habitat and distribution can be integral to such agreements, allowing foresight and the development of adaptive measures.

More generally, these results also highlight the emerging potential of marine-ecological forecasting as a climate change adaptation tool (IPCC, 2019). While we have focused here on the North Atlantic region, annual and multi-annual forecast skill is present in many other large marine ecosystems (Tommasi *et al.*, 2017) and can underpin the development of similar forecasts elsewhere. This technology is also particularly relevant to Small Island Developing States (SIDS) and the Global South, where ocean dependency and climate risk are amongst the highest in the world (Allison *et al.*, 2009; Barange *et al.*, 2014; Blasiak *et al.*, 2017). Regularly produced global-scale decadal-forecasts (Kushnir *et al.*, 2019) can support relevant climate services and thereby the sustainable development and climate adaptation of these nations (IPCC, 2019), for example via the UN Decade of Ocean Science for Sustainable Development, with it’s clear focus on “A Predicted Ocean”. Decadal-scale forecasts of the ocean, and of the life in it, thereby represent a tremendous opportunity for cutting edge climate science to have a direct benefit for the local communities, businesses and individuals that are most at risk from a changing and variable oceanic climate.

## Acknowledgements

The research leading to these results has received funding from the European Community’s Seventh Framework Programme (FP7 2007–2013) under grant agreement No 308299 (NACLIM) and the European Union’s Horizon 2020 research and innovation programme under grant agreement No 727852 (Blue-Action). The CESM project is supported primarily by the US National Science Foundation (NSF). The National Center for Atmospheric Research (NCAR) is a major facility sponsored by the US NSF under Cooperative Agreement 1852977. Computing and data storage resources for CESM-DPLE simulations, including the Cheyenne supercomputer (doi:10.5065/D6RX99HX), were provided by the Computational and Information Systems Laboratory (CISL) at NCAR. We acknowledge the World Climate Research Programme, which, through its Working Group on Coupled Modelling, coordinated and promoted CMIP6. We thank the climate modeling groups for producing and making available their model output, the Earth System Grid Federation (ESGF) for archiving the data and providing access, and the multiple funding agencies who support CMIP6 and ESGF.

## Author Contributions

MRP conceived the idea, performed the analyses and drafted the manuscript. AKM and MRP developed the blue whiting ecological niche model. NK, SGY, SY, GD and DM provided climate prediction model outputs and advised on their integration into the analysis. All authors contributed to revision of the manuscript and have approved the final version.

## Methods

### Study Region

We focus on the northern North Atlantic as the basis for this work. Decadal prediction experiments have shown this region to be one of the most predictable on the decadal scale (Kirtman *et al.*, 2013; Frölicher *et al.*, 2020). Studies have shown multi-annual to decadal predictability for sea surface temperature (Matei *et al.*, 2012b), upper ocean heat content (Yeager *et al.*, 2012), the Atlantic Meridional Overturning Circulation (AMOC; (Matei *et al.*, 2012a), CO_2_ uptake (Li *et al.*, 2016), and the dynamics of the North Atlantic subpolar gyre (Wouters *et al.*, 2013; Yeager and Robson, 2017; Yeager, 2020). This high underlying predictability of the physical system makes the North Atlantic an ideal candidate in which to develop decadal ecological forecasts and climate services (Payne *et al.*, 2017; Tommasi *et al.*, 2017).

### Fish Species and Ecological Niche Models (ENMs)

We focus on three fish species as case studies in the North Atlantic region (Extended Data Table 1); bluefin tuna (*Thunnus thynnus*), blue whiting (*Micromesistius poutassou*), and mackerel (*Scomber scombrus*), each of which has shown significant shifts in their spatial distribution in recent decades. For each of these species there is also an established body of knowledge and quantitative models that characterise the mechanisms underlying these shifts. These so-called ecological niche models (also known as species distribution models) parameterise the relationship between observations of the species and the physical environment, and are most commonly used to generate projections of habitat and distributional shifts of species under a changing climate. Their extension to predictive settings is therefore a natural one, and several applications on the near-real time and seasonal forecast scale are already established operationally (Hobday et al., 2011; Eveson et al., 2015; Payne et al., 2017; Hazen et al., 2018).

Recent shifts in the distribution of mackerel are amongst the most well-known examples of fish distributional shifts. The feeding distribution of mackerel expanded northwards and westwards to Iceland in 2007 (Astthorsson *et al.*, 2012) and Greenland in 2011 (Jansen *et al.*, 2016), leading to international conflicts over fishing rights on this species (Spijkers and Boonstra, 2017). A wide variety of explanations for these shifts can be found, with the effects of climate change and density-dependent expansion being the most common (Olafsdottir *et al.*, 2016; van der Kooij *et al.*, 2016; Nikolioudakis *et al.*, 2018). However, the distribution of mackerel is also clearly limited by temperature, with 8.5°C serving as a lower threshold (Jansen *et al.*, 2016; Nikolioudakis *et al.*, 2018). We therefore used the 8.5°C August-mean isotherm as a threshold for the habitat of this species. We focused on the waters around Greenland (specifically the exclusive economic zone south of 70°N) as a study region: being at the range limit of mackerel, this region is the most exposed to variations in the distribution of the species.

Bluefin tuna are pelagic top-predators that are widely distributed throughout the North Atlantic. The thermally suitable feeding habitat of this species expanded by 800 000 km^2^ from the mid-1980s to the early 2010s, leading to the first-ever observation of the species in Denmark Strait in 2012 (MacKenzie *et al.*, 2014). Unusually for fish, they have the ability to regulate their body temperature: their core body temperature is therefore often above the surrounding waters. Data-storage tags measuring both internal and external temperatures show that the species can dive into colder waters during the day for short periods to feed (e.g., horizontally across fronts or vertically across the thermocline), during which time the core temperature starts to drop, but then return to surface waters during the night to rest and warm-up again (Walli *et al.*, 2009). Such studies suggest that the species therefore needs access to surface waters of at least 10-11°C to support foraging, which can be interpreted as a natural limitation on the distribution of the species and definition of habitat (MacKenzie *et al.*, 2014). These conclusions are also seen in the results of empirical ecological niche models (Fromentin *et al.*, 2014; Druon *et al.*, 2016; Muhling *et al.*, 2017), that arrive at similar thresholds based on observations. Entirely independently, mechanistic bioenergetics modelling of the oxygen requirements and aerobic capacity of the species also reached a similar result (Muhling *et al.*, 2017). Like others (MacKenzie *et al.*, 2014; Jansen *et al.*, 2020), we therefore employ the 11°C isotherm for the August mean (the warmest month in the region) to define the maximum limit of thermally-suitable feeding habitat for this species in the northern North Atlantic.

Blue whiting is a small mesopelagic species found widely throughout the eastern North Atlantic. The species supports a large commercial fishery, primarily for industrial uses, that has varied greatly over time: in 2004 it was the world’s third largest fishery, with catches of 2.4 million tonnes (FAO, 2007). While smaller sub-populations exist, the largest stock, and the one that supports the majority of the fishery, migrates between its feeding grounds in the Norwegian Sea and spawning grounds to the west of Great Britain and Ireland in the Rockall Trough region. Spawning takes place from February to April (Pointin and Payne, 2014) and the spawning distribution varies substantially between years, expanding and contracting on and off the Rockall Plateau (Hátún *et al.*, 2009a). Initial work linked these changes to the large-scale dynamics of the North Atlantic sub-polar gyre (Hátún *et al.*, 2009b); however, more recent work has narrowed this view down to the local salinity conditions (Miesner and Payne, 2018) (which in turn is shaped by the basin-scale dynamics of the gyre). This work was based on approximately 34 000 observations (1100 presences) of blue whiting larvae in this region from 1951-2005 from the Continuous Plankton Recorder (CPR), which, in addition to the planktonic species for which it is best known, also regularly captures fish larvae (Corten and Lindley, 2003; Edwards *et al.*, 2011). An ecological niche model was developed and parameterized based on this data, using latitude, day of year, bathymetry, the solar elevation angle and environmental variables (averaged over 250-600 m) as predictors. The likelihood of observing blue whiting larvae in the CPR was found to have a dome-shaped response to salinity, with larvae occurrence limited to salinities between 35.3 and 35.5 psu. This model shows good agreement with independent observations from both scientific surveys and the fishery on the stock, and currently forms the basis of operational forecasts of the spawning distribution (ICES, 2018). The full ecological niche model (Miesner and Payne, 2018) was applied here to define suitable spawning habitat for this species, but was focused on the northern component (Pointin and Payne, 2014) where most of the variability has been observed.

### Physical Observations

Two different datasets were used as the basis for observations of the physical environment. Sea surface temperature estimates were based on the HadISST v1.1 product (Rayner *et al.*, 2003), while sub-surface salinity estimates were based on the EN4 product (v4.2.1 analysis, with Gouretski and Reseghetti (2010) corrections to the source profile data (Good *et al.*, 2013)), both from the UK Met Office. Both products are high-quality, internationally recognized estimates of the state of the ocean covering an extended time period (HadISST: from 1860, EN4 from 1900) and are presented on a regular 1° grid as monthly averages.

### Decadal Forecast Models

An ensemble of five decadal prediction systems was collated for this analysis: all models followed the CMIP6 Decadal Climate-Prediction Project (DCPP) protocol (Boer *et al.*, 2016). For each decadal prediction system, a database of retrospective forecasts was available based on annual initialisations. For each of these initialisations, a fully-coupled (ocean, sea ice, land, atmosphere) model was run freely from the given starting point to generate forecasts up to 10 years after the initialisation. Multiple realisations were available for each of the initialisations for each of the model systems, to give a grand ensemble of 85 members. Details are given in the relevant references (Extended Data Table 2).

### Uninitialized projections

We used uninitialized climate projections from the IPCC’s CMIP6 experiment (Eyring *et al.*, 2016) as one form of reference forecast. We selected a temporal subset of SST and salinity model outputs for the “historical” (covering the years 1960-2014) and “ssp585” (2015-2020) experiments. We used one realization from each model system, with the “Variant Label” identifier being maintained between the two experiments. Only models that fully covered the comparison period (1960-2018) were retained. Model outputs presented on sigma or density vertical axes or unstructured horizontal grids were excluded due to difficulties in incorporating them into the processing chain. Native model resolution (grid label “gn”) was used as the first preference, where available, followed by lower-resolution regridded products (“gr”,“gr1”). After this selection process, 35 models of salinity and 44 models of SST were incorporated into the analysis (Extended Data Table 3).

### Data Processing

Model and observational ocean (temperature and salinity) data were processed in the same manner. Data stored at multiple model levels (i.e., sub-surface salinity) were first extracted and then thickness-averaged over the 250-600-m depth range to produce two-dimensional fields for each time-step on the native model grid. The months of interest were then extracted from all fields and regridded using bilinear interpolation onto a common regular 0.5° latitude-longitude grid covering the region of interest.

### Observational Climatologies

Extracted and regridded observational data were used to generate monthly climatologies by grid-point averaging based on the 30-year period from 1985-2014 (inclusive). This period was chosen to cover the first 30 years for which predictions were used for all three fish case studies.

### Bias correction

Processed model outputs were corrected for bias. Climate model outputs often show systematic biases relative to observed values that are spatially variable. In the case of climate prediction systems, these biases can also vary as a function of forecast lead time due to the “forecast drift” phenomenon associated with adjustment from the initialized state to the model attractor (Magnusson *et al.*, 2013). Model outputs were bias-corrected following the “full field” approach, irrespective of the model initialisation technique applied (Choudhury *et al.*, 2017). Briefly, the climatological field of each variable (salinity and SST) was calculated for a given model and forecast lead time by grid-point averaging over the same 30 year period as the observational climatologies. The individual members of the forecast ensemble were then converted into a forecast anomaly based on this climatology for a given lead. Bias-adjusted full-field forecasts were then produced by adding the appropriate observational climatology to the forecast anomaly.

### Ensemble Means

Mean forecasts of environmental parameters for each model system were produced by averaging the forecast fields across the realisations for that forecast model. The mean across all 85 realisations in the ensemble was also calculated to produce a grand-ensemble mean forecast (“Grand Ens.”): in this way, each realisation was given equal weighting in the forecast, irrespective of the model system it came from or the number of “siblings” it may have. We also considered the mean-of-model-means approach, where each the mean-forecasts from each climate prediction were averaged, thereby giving each model system equal weight. However, the performance of this approach was indistinguishable from (or worse than) the grand ensemble approach, and is not presented here.

### Habitat Statistics

The skill of the near-term forecasts was evaluated based on retrospective forecasting (also known as hindcasting: the meaning of this term can differ between fields and so is avoided here). The ecological niche models (ENM) described above were applied to the bias-corrected full-field forecasts to generate a comparable set of retrospective habitat forecasts. Where the ENM gave a binary outcome (suitable / unsuitable habitat), the total area of suitable habitat was calculated directly. The blue whiting ENM (Miesner and Payne, 2018) however returned the probability of observing larvae and calibration is therefore needed prior to calculating the spawning habitat area for the observation of adult fish: the threshold was chosen to ensure agreement between the upper quartile of the annual adult distribution areas observed and the corresponding habitat estimates. The ENMs were also applied to observational datasets to generate observationally-based estimates of the area of habitat in a given year (“observed habitat” in Figures).

### Persistence and Binned Forecasts

Persistence forecasts used as the primary choice of reference forecast: a forecast is viewed as skillful if it can outperform such a baseline (Joliffe and Stephenson, 2012). Persistence forecasts were generated by propagating the habitat statistics calculated in a given year forward for up to a maximum of the 10-year forecast horizon considered here. Binned forecasts (i.e., the average over a multi-year period) were calculated based on 3, 5, 7, and 9-year windows of habitat statistics from all data sources (observations, persistence, uninitialized models and forecast models), and assigned a lead time corresponding to the centre of the mean window. The skill of both binned and persistence forecasts was then assessed in the same manner as for other data sources.

### Forecast Verification and Skill

Forecast skill was assessed by comparing the estimates of habitat based on observed environmental variables with forecasts of habitat based on the various forecast approaches. The retrospective forecast databases available differ in their length, and a common comparison period was chosen as 1961-2018 (inclusive) for SST-based variables, based on the intersection of available model-coverage. Initial explorations of salinity forecasts, however, revealed substantial inconsistencies between observational products prior to the mid-1980s that propagated into the initial conditions used to initialise the forecasts. In the absence of agreement between observational products about salinity in this region, we therefore limited the salinity comparison period to 1985-2018 (inclusive).

Forecast skill was quantified using multiple metrics including the Pearson correlation coefficient (*r*), the Mean Squared Error Skill Score (MSESS) and the Continuous Ranked Probability Skill Score (CRPSS) between “observed” and “predicted” habitat statistics (Joliffe and Stephenson, 2012). Skill-scores were calculated relative to the mean and standard deviation of the habitat statistics over the climatological period. CRPSS scores were calculated for the grand ensemble by considering the forecasts across all 85 ensemble members. Confidence intervals around each of these metrics were generated for each lead time by 1) pairwise resampling of years with replacement; 2) recalculating the appropriate metric; and 3) repeating the process 1000 times.

### Distribution and Abundance Data

Observations of distribution shifts suitable for verifying forecasts can be challenging to obtain: scientists rarely monitor a species in an area where it is not normally found. While we were unable to find suitable datasets for bluefin tuna and mackerel, the distribution of blue whiting has been the subject of routine scientific monitoring surveys since the early 1980s: since 2004 these surveys have been coordinated and standardised as the International Blue Whiting Spawning Stock Survey (ICES, 2015). Observations of blue whiting from this survey on a regular 2° × 1° grid were first used to identify the core 99% of the distribution in each year and the area occupied was then calculated. Estimates of the spawning stock biomass were obtained from the ICES Standard Graph Database (http://standardgraphs.ices.dk/) for the most recent stock assessment performed in 2020.

### Data availability

Climate predictions and projections analysed in this study are available from the CMIP data archives https://esgf-node.llnl.gov/projects/cmip6/. CESM-DPLE data are available from http://www.cesm.ucar.edu/projects/community-projects/DPLE/. HadISST data is available from https://www.metoffice.gov.uk/hadobs/hadisst/ and EN4 data from https://www.metoffice.gov.uk/hadobs/en4/

### Code availability

The code used during the current study is available from the corresponding author on reasonable request.

---

**Extended Data Fig. 1.**
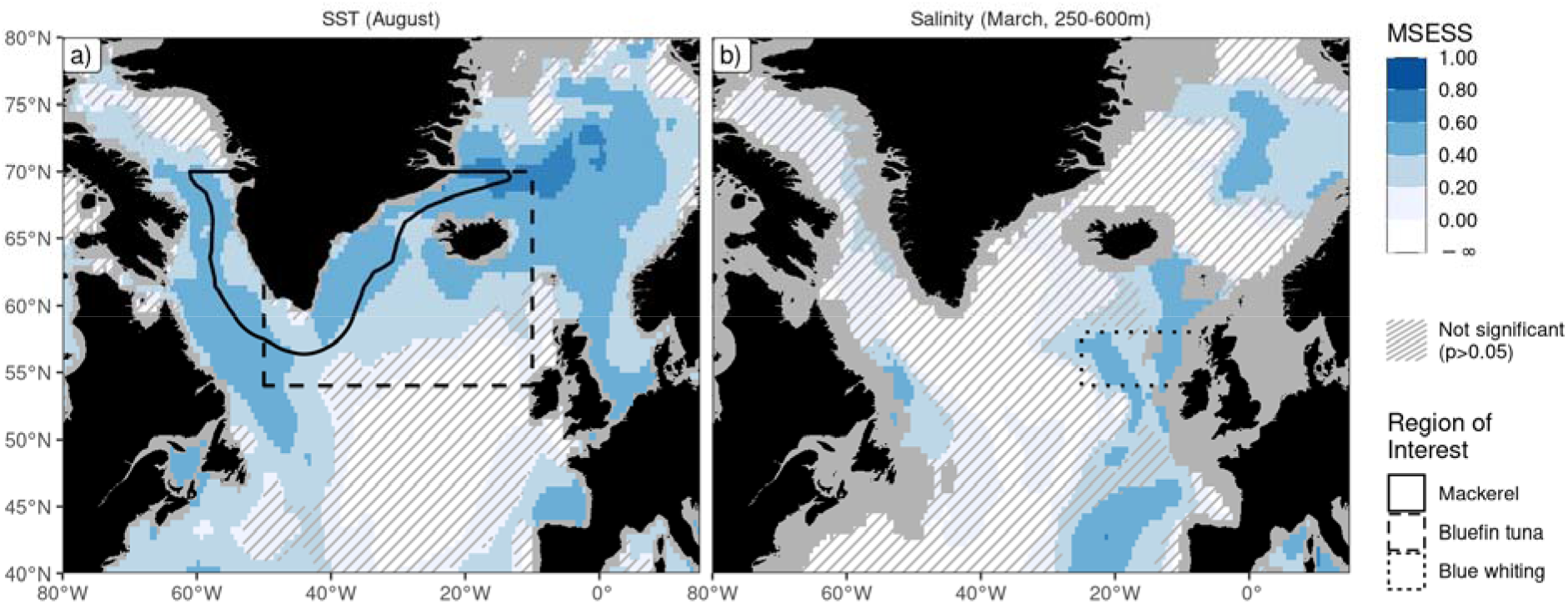
Absolute physical forecast skill. As for Fig.1 but showing mean squared-error skill score (MSESS) (rather than Pearson correlation) as a measure of forecast skill. Predictive skill of physical variables underlying ecological forecasts showing a) sea surface temperature (SST) in August and b) sub-surface (250-600 m) salinity in March with a lead time of five years. Each grid point is coloured according to the local MSESS estimate. Forecast skill is for the grand ensemble mean forecast, i.e., averaged across the individual realisations from all model systems. Regions where the MSESS is not significantly greater than 0 (at the 95% confidence level) are cross-hatched. Lines mark the polygons over which ecological forecasts are integrated in subsequent analyses.

**Extended Data Fig. 2.**
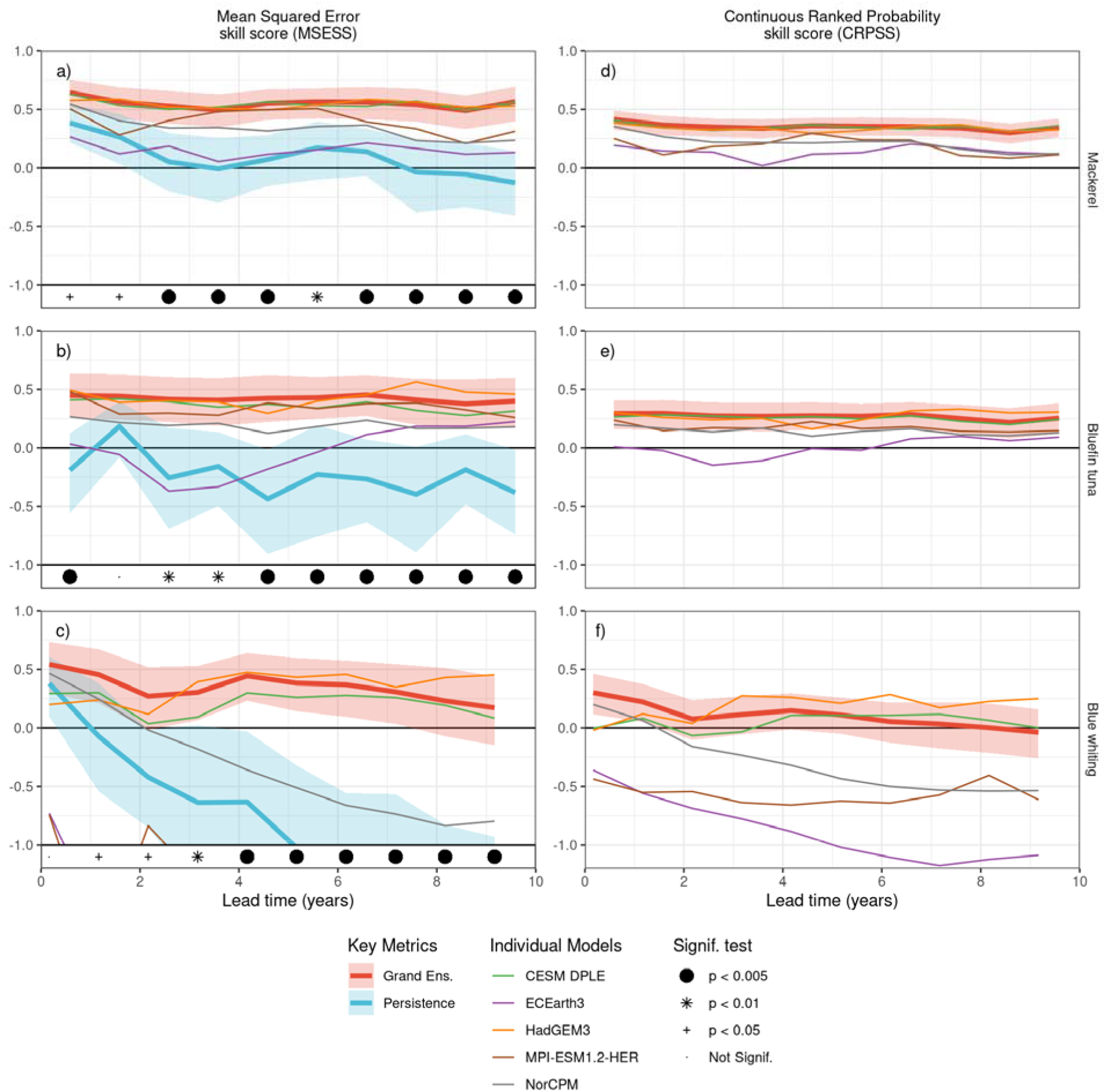
Additional metrics. As for Fig. 2 but showing additional metrics of forecast performance. The ability to correctly estimate the absolute habitat area is indicated by the Mean Squared Error skill score (MSESS) (panels a-c), while the Continuous Ranked Probability skill score (CRPSS) (panels d-f) indicates the probabilistic skill of the forecast distribution. Skill is shown for the habitat area of mackerel (panels a and d), bluefin tuna (b and e) and blue whiting (c and f). Skill metrics between the forecast and observed indicator values are plotted as a function of forecast lead-time into the future. Forecast skill is shown for the individual members of the model ensemble (light weighted lines) and for the grand-ensemble forecast (heavy red line). The skill of persistence forecasts (heavy blue lines) are also shown for reference where it can be defined (i.e. for MSESS): shaded areas for both these key metrics denote the 90% confidence interval. The hypothesis that the ensemble mean forecast outperforms persistence (i.e. a one-tailed test) is tested for each lead time, and denoted with symbols at the bottom of each panel. Both MSESS and CRPSS skill scores are calculated relative to the climatological statistics of each metric.

**Extended Data Fig. 3.**
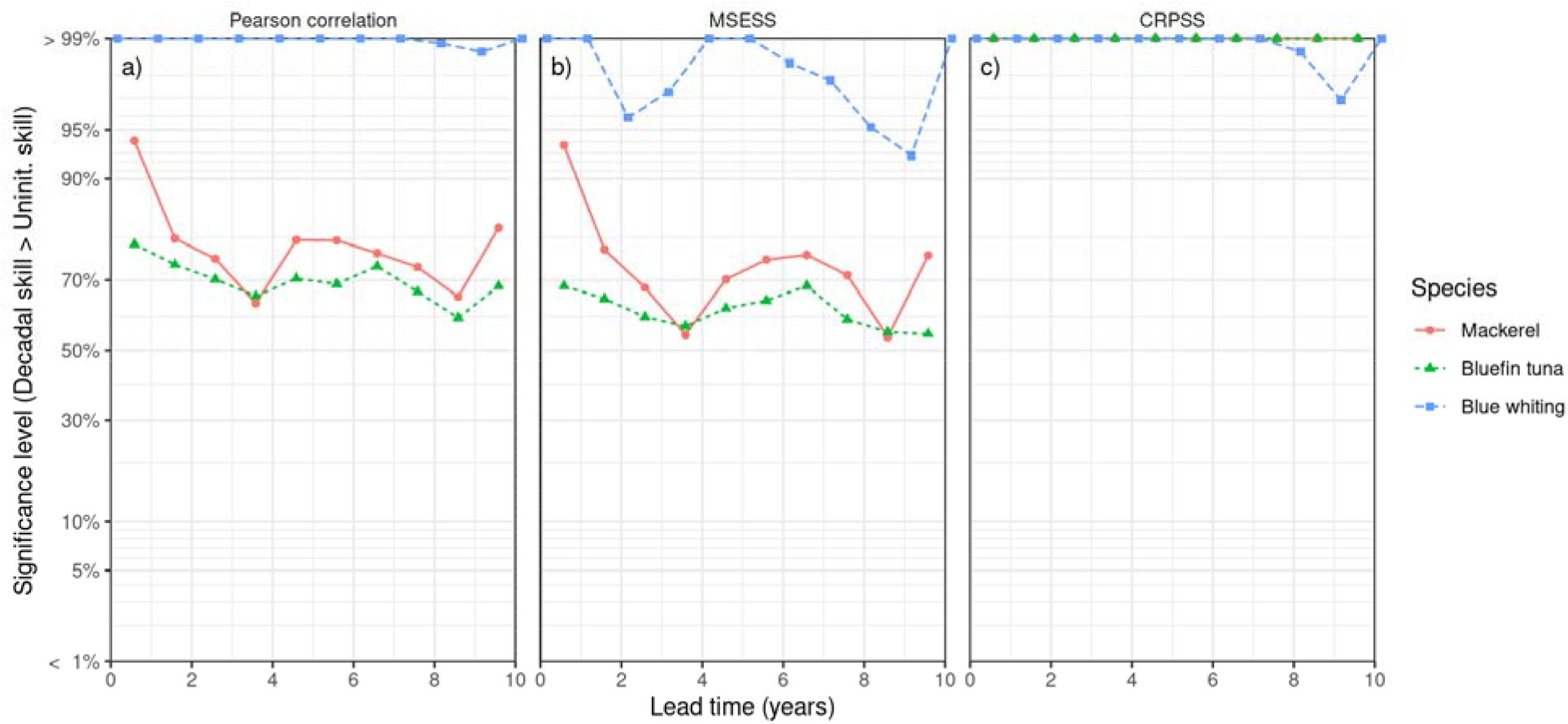
Significance levels against uninitialized forecasts. The significance of decadal forecast skill when compared against the skill of uninitialized forecasts (rather than persistence forecasts) for lead times of 0-10 years is shown for all species and for a) pearson correlation coefficient, b) the mean-squared error skill score (MSESS) and c) continuous ranked probability skill scores (CRPSS). Significance levels (1 - *p* values) are plotted on the vertical axis for a one-sided test that the given skill of the decadal forecast system is greater than the uninitialized skill. Note the non-linear (probit) scale on the vertical axis. Significance levels outside the axis ranges are plotted at the top or bottom of each panel.

**Extended Data Table 1.** Ecological niche models used in this study.

| Species | Region | Environmental Variable and Month of Interest | Ecological Niche Model (ENM) | References |
| --- | --- | --- | --- | --- |
| Mackerel (*Scomber scombrus*) | Greenlandic exclusive economic zone, south of 70°N | Sea surface temperature in warmest month (August) | Suitable habitat is warmer than 11°C | (Jansen *et al.*, 2016; Nikolioudakis *et al.*, 2018) |
| Bluefin tuna (*Thunnus thynnus*) | Irminger Sea, Denmark Strait and waters south of Iceland. | Sea surface temperature in warmest month (August) | Suitable habitat is warmer than 8.5°C | (MacKenzie *et al.*, 2014; Muhling *et al.*, 2017; Jansen *et al.*, 2020) |
| Blue whiting (*Micromesistius poutassou*) | Rockall Trough and Rockall Bank, west of Great Britain and Ireland | Salinity between 250 and 600 m depth (March) | Empirical model. Optimal salinity between 35.3 and 35.5 psu | (Hátún *et al.*, 2009b; Pointin and Payne, 2014; Miesner and Payne, 2018) |

**Extended Data Table 2.** Forecast systems and ensemble sizes used in this study.

| Forecast Centre | Model Name | Ocean Resolution | Start dates Ensemble size | References |
| --- | --- | --- | --- | --- |
| Bjerknes Center for Climate Research, Norway | NorCPM1 | Tripolar, 1° grid, 53 vertical levels on density coordinates | 1960-2018<br>20 members | (Bethke *et al.*, 2021) |
| Danish Meteorological Institute, Denmark | EC-Earth3 | Tripolar 1° grid with meridional refinement down to 1/3° in the tropics; 75 levels | 1960-2018<br>10 members | (Döscher *et al.*, 2021) |
| Max Planck Institute for Meteorology, Germany | MPI-ESM-1.2-HER | Tripolar, ~ 0.4° grid. 40 vertical levels. | 1960-2018<br>5 members | (Müller *et al.*, 2018; Mauritsen *et al.*, 2019) |
| Met Office Hadley Centre, UK. | HadGEM3-GC31-MM | Tripolar ~0.25° grid, 75 vertical levels | 1960-2018<br>10 members | (Williams *et al.*, 2018) |
| National Center for Atmospheric Research, USA | CESM DPLE | Nominal 1° horiz. with meridional refinement down to ~0.3° at the Equator; 60 vertical levels | 1955-2018<br>40 members | (Yeager *et al.*, 2018; Yeager, 2020) |

**Extended Data Table 3.**
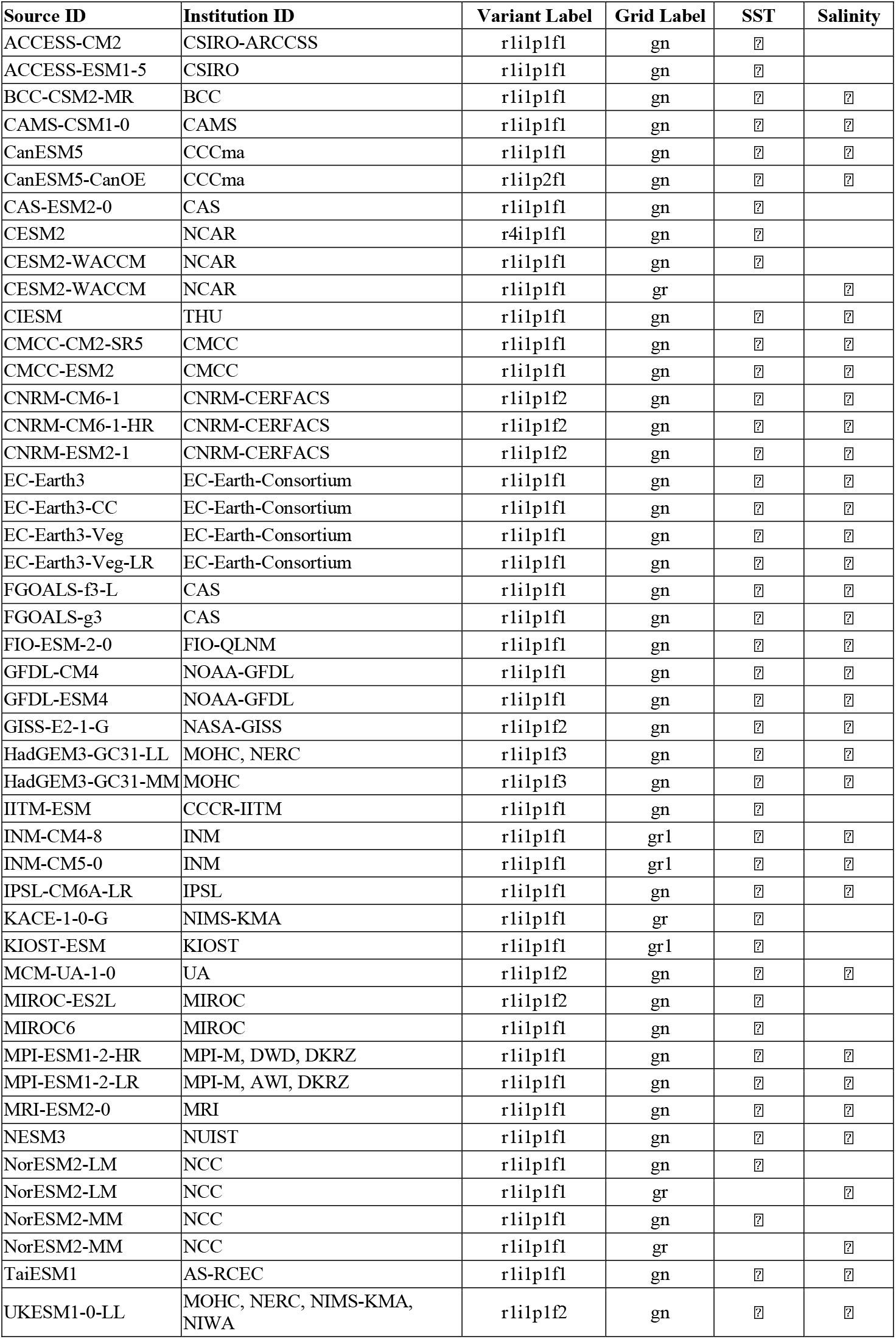
CMIP6 models and representative variants used as uninitialized models. Ticks indicate that the given model, variant and gridded product were used for uninitialized forecasts of either sea surface temperature (SST) or salinity. In total, 35 models were used for salinity and 44 for SST.

## References

Allison, E. H., Perry, A. L., Badjeck, M.-C., Neil Adger, W., Brown, K., Conway, D., Halls, A. S., et al. 2009. Vulnerability of national economies to the impacts of climate change on fisheries. Fish and Fisheries, 10: 173–196.

Astthorsson, O. S., Valdimarsson, H., Gudmundsdottir, A., and Oskarsson, G. J. 2012. Climate-related variations in the occurrence and distribution of mackerel (Scomber scombrus) in Icelandic waters. ICES Journal of Marine Science, 69: 1289–1297.

Barange, M., Merino, G., Blanchard, J. L., Scholtens, J., Harle, J., Allison, E. H., Allen, J. I., et al. 2014. Impacts of climate change on marine ecosystem production in societies dependent on fisheries. Nature Climate Change, 4: 211–216.

Bethke, I., Wang, Y., Counillon, F., Keenlyside, N., Kimmritz, M., Fransner, F., Samuelsen, A., et al. 2021. NorCPM1 and its contribution to CMIP6 DCPP. Geoscientific Model Development Discussions, 2021: 1–84.

Blasiak, R., Spijkers, J., Tokunaga, K., Pittman, J., Yagi, N., and Österblom, H. 2017. Climate change and marine fisheries: Least developed countries top global index of vulnerability. PLOS ONE, 12: e0179632.

Boer, G. J., Smith, D. M., Cassou, C., Doblas-Reyes, F. J., Danabasoglu, G., Kirtman, B., Kushnir, Y., et al. 2016. The Decadal Climate Prediction Project (DCPP) contribution to CMIP6. Geoscientific Model Development, 9: 3751–3777.

Bruno Soares, M., and Dessai, S. 2016. Barriers and enablers to the use of seasonal climate forecasts amongst organisations in Europe. Climatic Change, 137: 89–103.

Bruno Soares, M., Daly, M., and Dessai, S. 2018. Assessing the value of seasonal climate forecasts for decision-making. Wiley Interdisciplinary Reviews: Climate Change: 1–19.

Buontempo, C., and Hewitt, C. 2018. EUPORIAS and the development of climate services. Climate Services, 9: 1–4.

Choudhury, D., Sen Gupta, A., Sharma, A., Mehrotra, R., and Sivakumar, B. 2017. An Assessment of Drift Correction Alternatives for CMIP5 Decadal Predictions. Journal of Geophysical Research: Atmospheres, 122: 10282–10296.

Corten, A., and Lindley, J. A. 2003. The use of CPR data in fisheries research. Progress in Oceanography, 58: 285–300.

Döscher, R., Acosta, M., Alessandri, A., Anthoni, P., Arneth, A., Arsouze, T., Bergman, T., et al. 2021. The EC-Earth3 Earth System Model for the Climate Model Intercomparison Project 6. Geoscientific Model Development Discussions: 1–90.

Druon, J.-N., Fromentin, J.-M., Hanke, A. R., Arrizabalaga, H., Damalas, D., Tičina, V., Quílez-Badia, G., et al. 2016. Habitat suitability of the Atlantic bluefin tuna by size class: An ecological niche approach. Progress in Oceanography, 142: 30–46.

Edwards, M., Healouet, P., Halliday, N., Beaugrand, G., Fox, C., Johns, D. G., Licandro, P., et al. 2011. Fish larvae atlas of the NE Atlantic. Results from the Continuous Plankton Recorder survey 1948-2005. Sir Alister Hardy Foundation for Ocean Science, Plymouth, U.K. 22 pp.

Eyring, V., Bony, S., Meehl, G. A., Senior, C. A., Stevens, B., Stouffer, R. J., and Taylor, K. E. 2016. Overview of the Coupled Model Intercomparison Project Phase 6 (CMIP6) experimental design and organization. Geoscientific Model Development, 9: 1937–1958.

FAO. 2007. The state of world fisheries and aquaculture 2006. Rome. 180 pp.

Frölicher, T. L., Ramseyer, L., Raible, C. C., Rodgers, K. B., and Dunne, J. 2020. Potential predictability of marine ecosystem drivers. Biogeosciences, 17: 2061–2083. Copernicus GmbH.

Fromentin, J.-M., Reygondeau, G., Bonhommeau, S., and Beaugrand, G. 2014. Oceanographic changes and exploitation drive the spatio-temporal dynamics of Atlantic bluefin tuna (Thunnus thynnus). Fisheries Oceanography, 23: 147–156.

Golden, C. D., Allison, E. H., Cheung, W. W. L., Dey, M. M., Halpern, B. S., McCauley, D. J., Smith, M., et al. 2016. Nutrition: Fall in fish catch threatens human health. Nature, 534: 317–320.

Good, S. a., Martin, M. J., and Rayner, N. a. 2013. EN4: Quality controlled ocean temperature and salinity profiles and monthly objective analyses with uncertainty estimates. Journal of Geophysical Research: Oceans, 118: 6704–6716.

Guisan, A., and Zimmermann, N. E. 2000. Predictive habitat distribution models in ecology. Ecological Modelling, 135: 147–186.

Hátún, H., Payne, M. R., Beaugrand, G., Reid, P. C., Sandø, A. B., Drange, H., Hansen, B., et al. 2009a. Large bio-geographical shifts in the north-eastern Atlantic Ocean: From the subpolar gyre, via plankton, to blue whiting and pilot whales. Progress In Oceanography, 80: 149–162.

Hátún, H., Payne, M. R., and Jacobsen, J. A. 2009b. The North Atlantic subpolar gyre regulates the spawning distribution of blue whiting (Micromesistius poutassou). Canadian Journal of Fisheries and Aquatic Sciences, 66: 759–770.

Hewitt, C., Mason, S., and Walland, D. 2012. The Global Framework for Climate Services. Nature Climate Change, 2: 831–832. Nature Publishing Group.

ICES. 2015. Manual for International Pelagic Surveys (IPS). Series of ICES Survey Protocols SISP 9 – IPS. 92 pp.

ICES. 2018. Interim Report of the Working Group on Seasonal to Decadal Prediction of Marine Ecosystems (WGS2D), 27–31 August 2018, ICES Headquarters, Copenhagen, Denmark ICES CM 2018/EPDSG:22. 42 pp.

IPCC. 2019. IPCC Special Report on the Ocean and Cryosphere in a Changing Climate. In press pp.

Jansen, T., Post, Sø., Kristiansen, T., Óskarsson, G. J., Boje, J., MacKenzie, B. R., Broberg, M., et al. 2016. Ocean warming expands habitat of a rich natural resource and benefits a national economy. Ecological Applications, 26: 2021–2032.

Jansen, T., Nielsen, E. E., Rodríguez-Ezpeleta, N., Arrizabalaga, H., Post, S., and MacKenzie, B. R. 2020. Atlantic bluefin tuna (Thunnus thynnus) in Greenland – mixed-stock origin, diet, hydrographic conditions and repeated catches in this new fringe area. Canadian Journal of Fisheries and Aquatic Sciences: cjfas-2020-0156.

Joliffe, I. T., and Stephenson, D. B. 2012. Forecast Verification: a Practitioner’s Guide in Atmospheric Sciences. 292 pp.

Kirtman, B., Power, S., Adedoyin, J. A., Boer, G. J., Bojariu, R., Camilloni, I., Doblas-Reyes, F. J., et al. 2013. Near-term climate change: Projections and predictability.

Kushnir, Y., Scaife, A. A., Arritt, R., Balsamo, G., Boer, G., Doblas-Reyes, F., Hawkins, E., et al. 2019. Towards operational predictions of the near-term climate. Nature Climate Change, 9.

Li, H., Ilyina, T., Müller, W. A., and Sienz, F. 2016. Decadal predictions of the North Atlantic CO2 uptake. Nature Communications, 7: 11076.

MacKenzie, B. R., Payne, M. R., Boje, J., Høyer, J. L., and Siegstad, H. 2014. A cascade of warming impacts brings bluefin tuna to Greenland waters. Global Change Biology, 20: 2484–2491.

Magnusson, L., Alonso-Balmaseda, M., Corti, S., Molteni, F., and Stockdale, T. 2013. Evaluation of forecast strategies for seasonal and decadal forecasts in presence of systematic model errors. Climate Dynamics, 41: 2393–2409.

Matei, D., Baehr, J., Jungclaus, J. H., Haak, H., Muller, W. a., Marotzke, J., Vecchi, G. a, et al. 2012a. Multiyear Prediction of Monthly Mean Atlantic Meridional Overturning Circulation at 26.5 N. Science, 338: 604–604.

Matei, D., Pohlmann, H., Jungclaus, J., Müller, W., Haak, H., and Marotzke, J. 2012b. Two Tales of Initializing Decadal Climate Prediction Experiments with the ECHAM5/MPI-OM Model. Journal of Climate, 25: 8502–8523.

Mauritsen, T., Bader, J., Becker, T., Behrens, J., Bittner, M., Brokopf, R., Brovkin, V., et al. 2019. Developments in the MPI-M Earth System Model version 1.2 (MPI-ESM1.2) and Its Response to Increasing CO2. Journal of Advances in Modeling Earth Systems, 11: 998–1038.

Meehl, G. A., Goddard, L., Boer, G., Burgman, R., Branstator, G., Cassou, C., Corti, S., et al. 2014. Decadal climate prediction an update from the trenches. Bulletin of the American Meteorological Society, 95: 243–267.

Merryfield, W. J., Baehr, J., Batté, L., Becker, E. J., Butler, A. H., Coelho, C. A. S., Danabasoglu, G., et al. 2020. Current and Emerging Developments in Subseasonal to Decadal Prediction. Bulletin of the American Meteorological Society, 101: E869–E896. American Meteorological Society.

Miesner, A. K., and Payne, M. R. 2018. Oceanographic variability shapes the spawning distribution of blue whiting (Micromesistius poutassou). Fisheries Oceanography, 27: 623–638.

Mitchell, S. M., and Prins, B. C. 1999. Beyond territorial contiguity: Issues at stake in democratic militarized interstate disputes. International Studies Quarterly, 43: 169–183.

Muhling, B. A., Brill, R., Lamkin, J. T., Roffer, M. A., Lee, S. K., Liu, Y., and Muller-Karger, F. 2017. Projections of future habitat use by Atlantic bluefin tuna: Mechanistic vs. correlative distribution models. ICES Journal of Marine Science, 74: 698–716.

Müller, W. A., Jungclaus, J. H., Mauritsen, T., Baehr, J., Bittner, M., Budich, R., Bunzel, F., et al. 2018. A Higher-resolution Version of the Max Planck Institute Earth System Model (MPI-ESM1.2-HR). Journal of Advances in Modeling Earth Systems, 10: 1383–1413.

Nikolioudakis, N., Skaug, H. J., Olafsdottir, A. H., Jansen, T., Jacobsen, J. A., and Enberg, K. 2018. Drivers of the summer-distribution of Northeast Atlantic mackerel (Scomber scombrus) in the Nordic Seas from 2011 to 2017; a Bayesian hierarchical modelling approach. ICES Journal of Marine Science, 76: 530–548.

Olafsdottir, A. H., Slotte, A., Jacobsen, J. A., Oskarsson, G. J., Utne, K. R., and Nøttestad, L. 2016. Changes in weight-at-length and size-at-age of mature Northeast Atlantic mackerel (Scomber scombrus) from 1984 to 2013: effects of mackerel stock size and herring (Clupea harengus) stock size. ICES Journal of Marine Science: Journal du Conseil, 73: 1255–1265.

Palmer, T. N., Doblas-Reyes, F. J., Hagedorn, R., and Weisheimer, A. 2005. Probabilistic prediction of climate using multi-model ensembles: From basics to applications. Philosophical Transactions of the Royal Society B: Biological Sciences, 360: 1991–1998.

Payne, M. R., Hobday, A. J., MacKenzie, B. R., Tommasi, D., Dempsey, D. P., Fässler, S. M. M., Haynie, A. C., et al. 2017. Lessons from the First Generation of Marine Ecological Forecast Products. Frontiers in Marine Science, 4.

Pfaff, A., Broad, K., and Glantz, M. 1999. Who benefits from climate forecasts? Nature, 397: 645–646.

Pinsky, M. L., Reygondeau, G., Caddell, R., Palacios-Abrantes, J., Spijkers, J., and Cheung, W. W. L. 2018. Preparing ocean governance for species on the move. Science, 360: 1189–1191.

Pinsky, M. L., Eikeset, A. M., McCauley, D. J., Payne, J. L., and Sunday, J. M. 2019. Greater vulnerability to warming of marine versus terrestrial ectotherms. Nature. Springer US.

Pointin, F., and Payne, M. R. 2014. A Resolution to the Blue Whiting (Micromesistius poutassou) Population Paradox? PLoS ONE, 9: e106237.

Poloczanska, E. S., Burrows, M. T., Brown, C. J., Molinos, J. G., Halpern, B. S., Hoegh-Guldberg, O., Kappel, C. V., et al. 2016. Responses of marine organisms to climate change across oceans. Frontiers in Marine Science, 3: 1–21.

Rayner, N. A., Parker, D. E., Horton, E. B., Folland, C. K., Alexander, L. V., Rowell, D. P., Kent, E. C., et al. 2003. Global analyses of sea surface temperature, sea ice, and night marine air temperature since the late nineteenth century. Journal of Geophysical Research, 108: 4407.

Smith, D. M., Eade, R., Scaife, A. A., Caron, L.-P. P., Danabasoglu, G., DelSole, T. M., Delworth, T., et al. 2019. Robust skill of decadal climate predictions. npj Climate and Atmospheric Science, 2: 1–10.

Smith, D. M., Scaife, A. A., Eade, R., Athanasiadis, P., Bellucci, A., Bethke, I., Bilbao, R., et al. 2020. North Atlantic climate far more predictable than models imply. Nature, 583: 796–800. Springer US.

Spijkers, J., and Boonstra, W. J. 2017. Environmental change and social conflict: the northeast Atlantic mackerel dispute. Regional Environmental Change, 17: 1835–1851. Regional Environmental Change.

Spijkers, J., Singh, G., Blasiak, R., Morrison, T. H., Le Billon, P., and Österblom, H. 2019. Global patterns of fisheries conflict: Forty years of data. Global Environmental Change, 57. Elsevier Ltd.

Street, R. B. 2016. Towards a leading role on climate services in Europe: A research and innovation roadmap. Climate Services, 1: 2–5. Elsevier B.V.

Tommasi, D., Stock, C. A., Alexander, M. A., Yang, X., Rosati, A., and Vecchi, G. A. 2017. Multi-Annual Climate Predictions for Fisheries: An Assessment of Skill of Sea Surface Temperature Forecasts for Large Marine Ecosystems. Frontiers in Marine Science, 4: 1–13.

van der Kooij, J., Fässler, S. M. M. M., Stephens, D., Readdy, L., Scott, B. E., and Roel, B. A. 2016. Opportunistically recorded acoustic data support Northeast Atlantic mackerel expansion theory. ICES Journal of Marine Science: Journal du Conseil, 73: 1115–1126.

Walli, A., Teo, S. L. H., Boustany, A., Farwell, C. J., Williams, T., Dewar, H., Prince, E., et al. 2009. Seasonal Movements, Aggregations and Diving Behavior of Atlantic Bluefin Tuna (Thunnus thynnus) Revealed with Archival Tags. PLoS ONE, 4: e6151.

Williams, K. D., Copsey, D., Blockley, E. W., Bodas-Salcedo, A., Calvert, D., Comer, R., Davis, P., et al. 2018. The Met Office Global Coupled Model 3.0 and 3.1 (GC3.0 and GC3.1) Configurations. Journal of Advances in Modeling Earth Systems, 10: 357–380.

Wouters, B., Hazeleger, W., Drijfhout, S., van Oldenborgh, G. J. J., and Guemas, V. 2013. Multiyear predictability of the North Atlantic subpolar gyre. Geophysical Research Letters, 40: 3080–3084.

Yeager, S. G., Karspeck, A., Danabasoglu, G., Tribbia, J., and Teng, H. 2012. A Decadal Prediction Case Study: Late Twentieth-Century North Atlantic Ocean Heat Content. Journal of Climate, 25: 5173–5189.

Yeager, S. G., and Robson, J. I. 2017. Recent Progress in Understanding and Predicting Atlantic Decadal Climate Variability. Current Climate Change Reports, 3: 112–127. Current Climate Change Reports.

Yeager, S. G., Danabasoglu, G., Rosenbloom, N. A., Strand, W., Bates, S. C., Meehl, G. A., Karspeck, A. R., et al. 2018. Predicting near-term changes in the earth system: A large ensemble of initialized decadal prediction simulations using the community earth system model. Bulletin of the American Meteorological Society, 99: 1867–1886. American Meteorological Society.

Yeager, S. G. 2020. The abyssal origins of North Atlantic decadal predictability. Climate Dynamics, 55: 2253–2271. Springer Berlin Heidelberg.

